# The hepcidin-ferroportin axis influences mitochondrial function, proliferation, and migration in pulmonary artery endothelial and smooth muscle cells

**DOI:** 10.1101/2023.10.09.561497

**Authors:** Theo Issitt, Quezia K Toe, Sofia L Pedersen, Thomas Shackshaft, Maziah Mohd Ghazaly, Laura West, Nadine D Arnold, Abdul Mahomed, George W Kagugube, Latha Ramakrishnan, Allan Lawrie, S John Wort, Gregory J Quinlan

**Affiliations:** NHLI, Faculty of Medicine, Imperial College London SW3 6LY UK; Univerity of Malaysia Terengganu, Malaysia; Department of Infection, Immunity & Cardiovascular Disease, University of Sheffield, S10 2RX, Sheffield, UK

**Keywords:** Hepcidin ferroportin axis, iron, IL-6, pulmonary artery endothelial cells, pulmonary artery smooth muscle cells, migration and proliferation, mitochondria, BMPRII

## Abstract

**Rationale:** Elevated circulating hepcidin levels have been reported in patients with pulmonary artery hypertension (PAH). Hepcidin has been shown to promote proliferation of human pulmonary artery smooth muscle cells (PASMCs) in vitro, suggesting a potential role in PAH pathogenesis. However, the role of human pulmonary artery endothelial cells (PAECs) as either a source of hepcidin, or the effect of hepcidin on PAECfunction has not previously been described.

**Objective:** To define the role of the hepcidin-ferroportin axis on the phenotype of pulmonary artery endothelial cells

**Methods and results:** PAECs treated with hepcidin, or IL-6 were investigated for both ferroportin and hepcidin release and regulation with immunofluorescence, mRNA levels and cellular release assays. Effects of hepcidin on PASMC and PAEC mitochondrial function was investigated using immunofluorescence and seahorse assay. Migration and proliferation of PASMC treated with conditioned media from hPAEC treated with hepcidin was investigated using the Xcelligence system and other tools.

PAECs express ferroportin; hepcidin treatment of PAECs results in mitochondrial iron accumulation and intracellular hepcidin biosynthesis and release. Conditioned media from hepcidin treated PAECs causes PASMCs to down regulate ferroportin expression whilst promoting migration and proliferation. Inhibition of hepcidin in PAEC conditioned media limits these responses. PASMC cellular and mitochondrial iron retention are associated with migratory and proliferative responses.

**Conclusions:** The Hepcidin-ferroportin axis is present and operational in PAECs. Modulation of this axis shows distinct differences in responses seen between PAECS and PASMCs. Stimulation of this axis in PAECS with hepcidin may well institute proliferative and migratory responses in PASMCs of relevance to pathogenesis of PAH offering a potential therapeutic target.

## Introduction

Pulmonary artery hypertension (PAH) is a rare, life-limiting disease with limited treatments options. Remodelling of resistance, pre-capillary, pulmonary blood vessels including sustained vasocontriction followed by the proliferation of pulmonary artery smooth muscle cells (PASMCs) and endothelial (PAECs) results in vessel muscularisation, lumen restriction and eventually obliterative lesions. The subsequent elevation in pulmonary vascular resistance leads to right sided heart failure and early death (Condon et al., 2019).

Several genes and genetic abnormalities have been identified in the heritable form of PAH with mutations in the bone morphogenetic protein receptor (BMPR) II the most common (Dannewitz Prosseda et al., 2020, Graf et al., 2018). Moreover, deficiency in BMPRII protein levels (and associated signalling molecules) have been shown in non-genetic and other forms of PAH cases (Machado et al., 2015), suggesting commonality, and potentially offering some mechanistic insight. A role for other members of the TGF-β super family of receptors has also been implicated in PAH (Rabinovitch, 2012).

Signalling molecules in the TGF-β superfamily including BMP/SMAD pathways are crucial for cell turnover and maintaining the balance between proliferation and apoptosis. In addition BMP /SMAD pathways are involved with numerous regulatory functions including that of iron homeostasis; specifically, through the control of the expression and release of the small peptide hormone, hepcidin (Finberg, 2013), the master regulator of body iron turnover and control. In this regard, disrupted iron homeostatic control has been implicated in the pathogenesis of idiopathic PAH (Soon et al., 2011) and related conditions, such as Eisenmenger syndrome (Van De Bruaene et al., 2011). Hepcidin functions via interaction and down-regulation of ferroportin activity, ferroportin being the only known cellular iron exporter in mammalian systems (Drakesmith et al., 2015). Ferroportin expression is largely restricted to expression in hepatocytes, erythroblasts, enterocytes and sub-types of macrophages, all cells with key function(s) for the iron regulation and reprocessing. However, ferroportin is also expressed more widely (Drakesmith et al., 2015, Ginzburg, 2019) including in human PASMCs where hepcidin treatment stimulates proliferation and cellular iron accumulation. Moreover, stabilisation of ferroportin membrane localisation and activity was able to reverse this proliferative response and iron accumulation despite the presence of hepcidin. (Ramakrishnan et al., 2018). Corroboration for the operation of the hepcidin-ferroportin axis in PASMCs of relevance to PAH has recently been provided *in vivo* and in human cells (Lakhal-Littleton et al., 2019), suggesting a localised iron regulatory axis may be operational in this vascular bed which may have implications for PASMC remodelling should homeostatic control be lost. Indeed, iron is a mitogen and co-factor for ribonucleotide reductase (Cotruvo and Stubbe, 2011) a key enzyme for DNA biosynthesis, and additionally for several cell cycle cyclins (Yu et al., 2007).

Therefore, implications for altered iron regulation in any such cells will likely profoundly impact mitochondrial function given the key requirement of iron for respiration and biosynthesis within these organelles. Moreover, cellular iron-loading is known to cause mitochondrial dysfunction enhancing intracellular reactive oxygen species (ROS) production, affecting intracellular cell signalling responses and ultimately cell fate (Rouault et al. 2016). Throughout the literature, a role for mitochondrial dysfunction in PAH is emerging (Zhang et al., 2022).

This study investigates the hepcidin-ferroportin axis in PAECs and PASMCs. We show that hepcidin exposure to PAECs and PASMCs causes strikingly different mitochondrial responses between cell types with PASMCs showing pronounced mitochondrial dysfunction. In addition, hepcidin exposure to PASMCs promotes enhanced hepcidin production and release from these cells, and further that these processes stimulate both migration and proliferation in PASMCs, all such findings which may be of relevance to vascular remodelling in PAH.

## Methods

### Tissue and cell culture

Human pulmonary artery smooth muscle cells (PASMC) were isolated and maintained as previously described using Dulbecco’s Modified Eagle’s Medium (DMEM) supplemented with 15% fetal calf serum (FCS), 2 mM glutamine, 100 U/ml penicillin, and 100 μg/ml streptomycin (Ramakrishnan et al. 2018). All human pulmonary artery endothelial cells (PAEC) were supplied fully characterised by Promocell and grown in endothelial cell basal medium 2 (ECM), with supplements and 2% fetal calf serum (Promocell). All cells were grown at 37°C in a humidified atmospheric incubator with 5% CO2.

### Treatments and conditioned media preparation

Both PAEC and PASMC were grown to a 90% confluent monolayer before trypsinisation with detach kit (Promocell) as per manufacturer’s instructions. Prior to treatments, PAEC or PASMC were starved overnight in serum restricted media (0.2% FCS containing ECM with supplements as supplied, excluding heparin and hydrocortisone or 0.2% FCS containing DMEM with 2 mM glutamine, 100 U/ml penicillin, and 100 μg/ml streptomycin). Treatment of cells with respective molecules was performed in serum restricted media. Hepcidin-25 was purchased from Peptides International and IL-6 from R&D Systems.

For production of conditioned media, PAECs were seeded in 12-well plates at a density of 7.5 x 10^4^ cells/well. Cultured cells were only used for experiments between passage 4-7. After starvation, cells were treated were serum restricted media (control), Hepcidin at 1 µg/mL or IL-6 at 10 ng/mL (unless otherwise specified). Following 24 hours of exposure, media was collected and used immediately for direct treatment upon cells or flash frozen in dry ice and kept at -80°C.

### Transfection

PAEC were transfected with lipofectamine RNAiMAX (Thermofisher, UK) transfecting agent with SMARTpool siRNA targeting HAMP (Dharmacon, USA cat#: L-014014-00-0005) or control ON-TARGETplus non-targeting control pool (Dharmacon, USA Cat#: D-001810-10-05)

### Gene expression

mRNA was collected from cultured cells using the RNAeasy system (Qiagen, UK) and converted to cDNA using the first strand cDNA synthesis kit (Thermofisher, UK) or iScript cDNA kit (Bio-Rad, UK) for total RNA synthesis. Amplification was performed with SYBR green reagents (Bio-Rad, UK).

### exCELLigence assay

In proliferation experiments with direct treatments, PASMCs were plated and starved as described in section 2.3. After 24 hours, cells were treated with 0.2% FCS DMEM and no further additives (control) for 90 minutes at 37°C. After blocking, the PASMCs were treated with 15% FCS DMEM and no further additives (control), or hepcidin at 1 µg/mL or 100 ng/mL, or IL-6 at 10 ng/mL or 1 ng/mL. Repeat treatments were performed 48 hours later.

Similar to the proliferation assay, the xCELLigence RTCA system can be used to measure cell migration in real time. The migration plates consist of wells separated into two connected chambers: cells are placed in the upper one, while conditioned media or other treatments are placed in the lower one. Between the two chambers, a microporous membrane allows cell migration. Migration is measured by the RTCA system as changes in electrical impedance; these occur as cells migrate and attach to the underside of the microporous membrane, which is covered in a gold microelectrode array (Figure X).

160 µL CM collected from PAECs (described in 2.2.) was added to the lower chambers, as well as ECM and 15% FCS DMEM controls. The top chamber was then affixed to the lower chamber and 50 µL of 0.2% FCS DMEM added to each upper chamber. The apparatus was then incubated at 37°C for 1 hour, after which a background reading of electrical impedance was taken. 100 µL PASMC solution was added to each upper chamber at a density of 15,000 cells/well. The plates were incubated at room temperature for 30 minutes to allow the cells to sediment before commencing RTCA Cell Index measurements every 15 minutes for 16 hours.

### BRDU

Cells were plated on 96 well plate (2500 cells/well). Cells were plated at 70% confluence, after initial adherence, the cells were serum starved (0.2% Serum) for 24h prior to treatment. Following treatment for 24h, BrdU was introduced (at the manufacturer recommended concentration) for an additional 24h incubation before harvesting the plates. Proliferation was quantified using anti-BrdU-POD antibody, according to the manufacturer’s instructions. All treatments were undertaken in triplicate.

### MTS assays

Cell proliferation was analysed by the CellTiter 96 ® Aqueous One Solution Cell proliferation Assay (Promega, UK). Following cell treatments after incubation period, the supernatant of each each (96-well plate) was removed and replace with 100 μL of fresh complete medium and 20 μL of MTS assay was added to each well, 3 blank wells were also added, this well only contained media and MTS assay only.

### Immunoblotting

Cells were lysed in RIPA buffer (Sigma-Aldrich, UK) with protease and phosphatase inhibitor cocktails (Sigma-Aldrich, UK) for 10 mins on ice. 40µg of protein was resuspended in laemelli buffer denatured at 95°C for 5mins, separated by SDS-PAGE and transferred to a nitrocellulose membrane. Blots were incubated over night at 4°C with primary antibodies for ferroportin (rabbit anti-FPN (Sigma-Aldrich, UK)) and α-tubulin (mouse anti-α-tubulin, Proteintech, UK). HRP-conjugated antibodies were used to visualise blots.

### Immunoflourescence

For experiments with fixed samples, PAECs and PASMCs were fixed in 4% paraformaldehyde in PBS for 7 min, permeabilised in 0.25% Triton X-100 in PBS for 5 min, blocked for 1 hour in 0.1% BSA in PBS and incubated overnight with the following primary antibodies: Rabbit anti-TOM20, (Proteintech, UK), DAPI and rabbit anti-FPN. The following conjugated secondary antibodies were used: Alexa-488 conjugated goat anti-rabbit/anti-mouse, Alexa-594 donkey anti-rabbit/anti-mouse or Alexa-700 donkey anti-rabbit secondary antibodies (ThermoFisher Scientific, UK) and DAPI (Sigma-Aldrich, UK). Images and z-stacks were acquired with a plan apochromat 40X oil objective on an SP8 inverted confocal microscope (Leica, UK). Maximal intensity projections were generated with ImageJ (NIH, Bethesda, USA) and normalized to DAPI pixel intensity.

### Mitotracker staining

PAECs and PASMCs were incubated with serum-free ECM or DMEM media respectively containing 125nM MitoTracker™ Orange CMTMros (ThermoFisher, UK) for 25 minutes. Cells were washed three times in PBS (Sigma-Aldrich, UK) and then processed as fixed samples as described. Mitochondrial area and volume per cell were calculated using Volocity (Perkin-Elmer, USA) through Z-stack analysis and thresholding. Nuclear volume was determined in Volocity by DAPI staining and used as a normalizing factor.

### Mito-FerroGreen live-staining

PAECs grown on imaging slides (Ibidi, Germany) were washed three times with HBSS supplemented with Ca^2+^ and Mg^2+^ (Sigma-Aldrich, UK) and incubated for 30 minutes with 5 µM Mito-FerroGreen and Hoescht 3342 (Sigma-Aldrich,UK). Cells were then washed three times with HBSS supplemented with Ca^2+^ and Mg^2+^ and live-imaged with SP8 inverted confocal microscope microscope with a plan apochromat 40X oil objective (Leica, UK). Mito-FerroGreen pixels were determined with ImageJ (NIH, Bethesda, USA).

### Mitosox live-staining

ECs were incubated with 5 µM Mitosox (Sigma-Aldrich, UK) and 5 µg/ml Hoescht 33342 (Sigma-Aldrich, UK) in HBSS with Ca^2+^ and Mg^2+^ (Life Technology, UK) for 8 minutes, washed three times with HBSS with Ca^2+^ and Mg^2+^ before imaging with a LSM780 confocal microscope with a plan apochromat 63X 1.4 NA oil objective (Zeiss, Germany). In some cases, cells were treated with 100 µM Deferoxamine (Sigma-Aldrich, UK) for 24 hours before imaging. Mitosox and Hoescht 33342 integrated density was determined with ImageJ (NIH, Bethesda, USA).

### Immunoflourescence analysis

Mitotracker CMTMros was measured to determine mitochondrial membrane potential. Its entry into mitochondrial is dependent upon the membrane potential and reduced fluorescence can indicate a reduction of this potential (Lin et al., 2013). Mitochondria retain these dyes regardless of flucuation of MMP and so variation over time cannot be measured (Kholmukhamedov et al., 2013, Poot et al., 1996). We have used this quantification here as an immediate measurement only. Cells were starved overnight and treated with hepcidin for 24hours. Then incubated with Mitotracker for 20 mins at 37C. Cells were then fixed and stained. Confocal z-stacks were compiled as z projections according to pixel max intensity and measurements taken in Volocity (Perkin-Elmer, USA). Fold change was calculated for average integrated density per cell relative to the control of that experiment n=3.

### Cell cycle analysis

Cell cycle was investigated using propidium iodide assay via flow cytometry following manufacturer’s instructions (Invitrogen). Briefly, cells were washed in PBS, trypsinised, pelleted and resuspended in PBS with staining solution, incubated for 30 mins in the dark and analysed with flow cytometer (BD biosciences).

### Gene expression qPCR

Total RNA was extracted form cells using the RNAeasy Mini preparation Kit (Qiagen, UK). Total RNA was measured by Nano-drop spectrophotometer 0.1-0.5 μg of total RNA was used to produced cDNA, using MLV reverse transcriptase (Invitrogen), 10 mM dNTPs, oligo-dT primers (Invitrogen) and RNase inhibitor (Applied Biosciences). Real-time PCR using SYBR green (Sensi-FAST lo-ROX, Meridian Bioscience) was carried-out on a Rotor-Gene 6000 PCR machine. The change in expression was normalised to control untreated samples.

### Statistics

Statistical analysis was performed on GraphPad (Prism). Graphs generated using GraphPad present data ± SEM (standard error of the mean) of the specified number of independent experiments.

## Results

This research has demonstrated presence of the hepcidin:ferroportin axis in PAEC and PASMC. Treatment of PAEC with hepcidin induced production of hepciding, IL-6 and other cytokines. Direct treatment of PAECs and PASMCs with hepcidin alone generated contrasting mitochondrial stress, whereby PAECs exhibited less mitochondrial stress than PASMC. Treatment of PASMC with media from PAECs treated with hepcidin induced proliferative, migratory and metabolic changes.

### Ferroportin and hepcidin are expressed and regulated in PAEC

Immunofluorescence revealed ferroportin distribution throughout hPAECs (Fig1A, control image). A reduction in ferroportin fluorescence following hepcidin treatment for 24 hours was observed indicating effective control of this regulatory axis by hepcidin (Fig1A, hepcidin image). Three hour and 6-hour hepcidin challenge in PAEC revealed a slight increase in ferroportin (Fig S1A). IL-6 challenge, a known regulator of hepcidin expression (Lee et al., 2005, Wrighting and Andrews, 2006) was ineffective for the control of ferroportin expression over this time frame (Fig 1A,IL-6 image). Western blot of hPAEC control lysates revealed a band around 50 kD, as expected for ferroportin, which we have reported previously (Toe et al., 2022). Basal levels of ferroportin mRNA were detectable in PAEC and expression was significantly reduced following 2 hours of stimulation with 10 ng/mL IL-6 and 1 µg/mL hepcidin (Toe et al., 2022).

**Figure 1.**
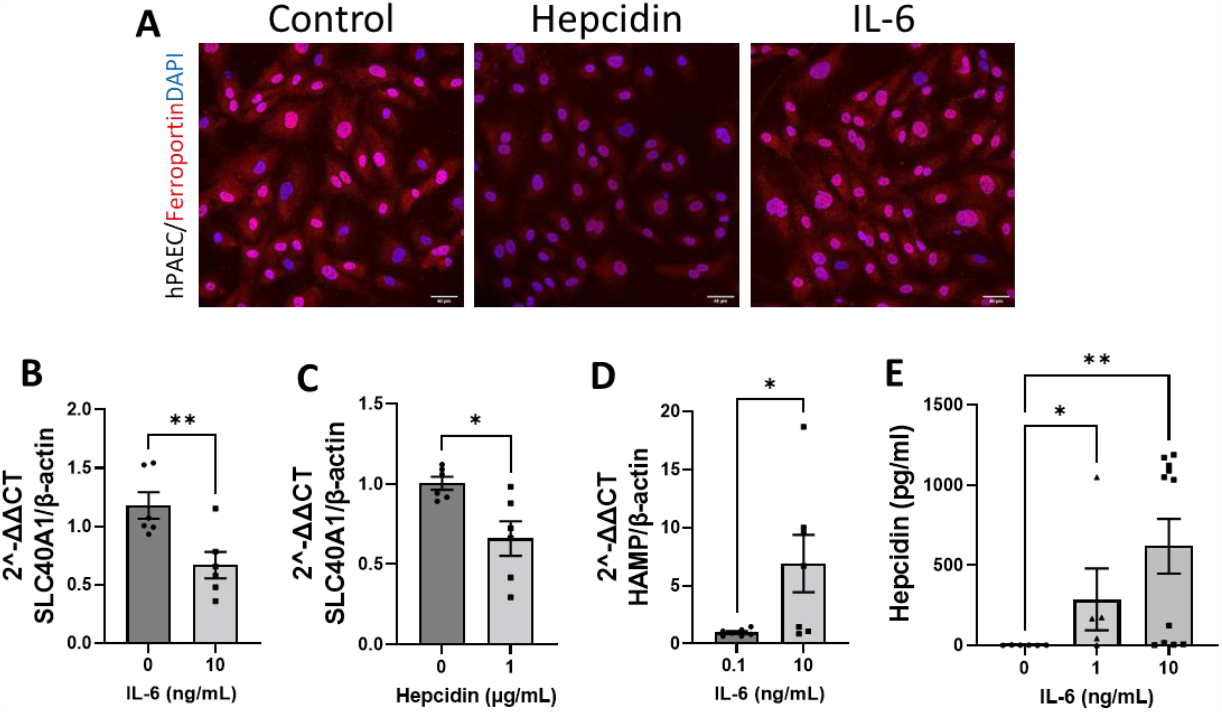
Ferroportin and Hepcidin expression and regulation in hPAEC. **(A)** Confocal images of hPAEC grown with normal media (control) or treated with 1 µg/mL hepcidin for 24 h or 10 ng/ml IL-6 and stained for ferroportin (red) and DAPI (blue), Scale bar 40pm. **(B)** Immunoblot of hPAEC lysates probed for TPN, α-tubulin shown as control. **(C, D)** Quantification of ferroportin (SLC40A1) mRNA by RT-qPCR in hPAECs treated with 10 ng/ml IL-6 (C) and 1 µg/mL hepcidin (D) for 2 hr, expressed as fold change of control (mean ± SEM; n = 5),; Student’s t test; ^**^p<0.01; ^*^p<0.05. **(E)** Hepcidin mRNA transcription in hPAECs after IL-6 stimulation for 2 hours. Data shown are mean ±SEM n=6. Unpaired t-test was performed. ^*^p<0.05. **(F)** Quantification of ELISA specific to Hepcidin for media supernatants of hPAEC untreated (control) or with 1 – l0ng/ml IL-6 for 24 hours (mean ± SEM; n - 5-12) Kruskal-Wallis test followed by Dunn’s post hoc test was performed; ^*^p< 0.05.

IL-6 challenge to PASMC was previously shown to cause these cells to produce and release hepcidin (Ramakrishnan et al., 2018). IL-6 treatment of hPAECs similarly revealed significantly reduced ferroportin mRNA (Fig 1B), similar to the response seen following hepcidin treatment (Fig 1C). IL-6 treatment of hPAEC also induced hepcidin mRNA (HAMP) at 1 hour (Fig 1D). Additionally, an increased cellular release of hepcidin protein was detectable in the media from PAECs treated with IL-6 for 24 hours (Fig 1E). Taken together these results effectively demonstrate the capacity for PAECs to produce both ferroportin and hepcidin and indicate a localised iron regulatory axis in these vascular cells.

Immunoflourescence of PASMCs following treatment with hepcidin also revealed a decrease in ferroportin after 24hours (Fig S1B). 24 hours IL-6 treatment of PASMC caused no change in ferroportin expression (Fig S1B). In line with observations in PAECs, mRNA for ferroportin increased initially, at 1 hour post treatment, but was ultimately reduced relative to control following hepcidin treatment (Fig S1C).

### Hepcidin treatment of hPAEC induces production of IL-6, and augmented hepcidin release

Hepcidin treatment of PAECs resulted in elevated levels of IL-6 mRNA in comparison to untreated controls and within 1 hour of exposure (Fig 2A). In addition, a significant increase of IL-6 protein release from PAECs was found in media from hepcidin treated PAECs by 24 hours, as determined by ELISA (Fig 2B). The ability of hepcidin to induce IL-6 expression has previously been demonstrated in PASMCs by our group (Ramakrishnan et al., 2018); these studies demonstrate a similar capacity for PAECs.

**Figure 2.**
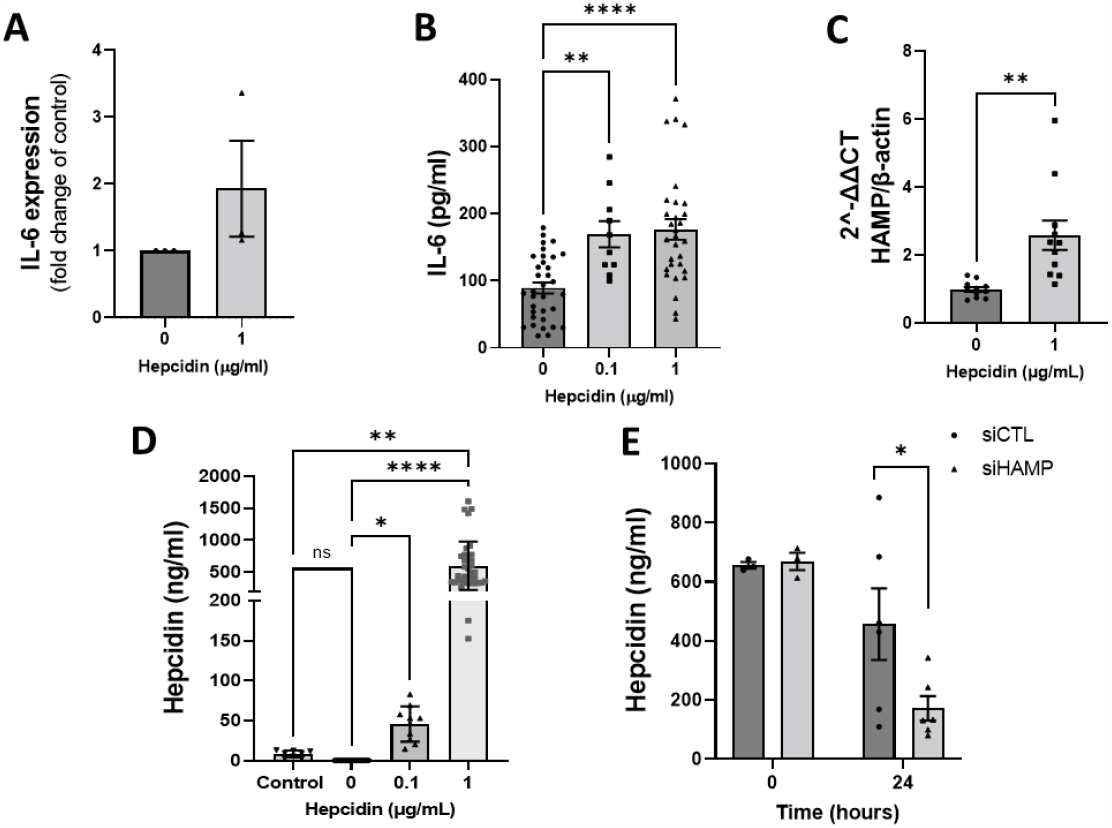
Hepcidin induces production of IL-6 and Hepcidin in hPAEC. hPAECs release IL-6 after hepcidin treatment **(A)**. IL-6 mRNA transcription in hPAECs after hepcidin stimulation for 5 hours. Data shown are mean +SEM n=3. (B) IL-6 release into hPAECs supernatants after 24 hours stimulation with hepcidin (0.1, 1 µg/mL). hPAECs supernatants were analysed by ELISA as described in section 2.6. Data shown are mean ±SEM n=12-36. **(C)** hepcidin mRNA transcription from hPAECs after treatment with hepcidin tor 1 hours. Data shown are mean ±SEM n=6. **(D)** ELISA analysis ot hepcidin in media supernatants of hPAEC untreated or with 0.1 or 1µg/ml Hepcidin for 24 hours. Control denotes media alone with 1pg/ml Hepcidin for 24 hours. Data shown are mean ±SEM n=8-30 **(E)** ELISA analysis of hepcidin In supernatants of hPAECs transfected with si-control or SI-HAMP treated with lpg/ml Hepcidin at time 0 (six hours after transfection) and 24 hours after transfection . Data shown are mean ± SEM; n = 3, 6. Unpaired t-test **(A, C, E)** or Kruskal-Wallis test followed by Dunn’s post hoc test was performed. **(B, D)**; ^****^p <0.0001; ^**^p < 0.01; ^*^p < 0.05.

Interestingly, treatment of PAECs with hepcidin was found to induce augmented release of hepcidin by these cells. Treatment with hepcidin revealed, firstly, a significant increase in HAMP mRNA at 1 hour (Fig 2C). This was followed with significant release of hepcidin into the media compared to control over 24 h, as determined by ELISA (Fig 2D). Media alone showed presence of low levels of hepcidin relative to that seen with cells. 0.1 and 1 µg/mL hepcidin treatment both induced significant hepcidin release by cells into the media (Fig 2D). Furthermore, this response was shown to increase over time, up to 24 h (Fig S2A). Comparison between cells incubated with hepcidin and cell only controls reinforced the role of hepcidin as mediator for subsequent hepcidin release from PAECs (Fig 2D). Downregulation of hepcidin (HAMP) through transfection of PAEC with targeting siRNA revealed a significant reduction in hepcidin present in the media of hepcidin treated cells compared with a control siRNA (Fig 2E).

Interestingly, hepcidin treatment in PAEC caused significantly increased release of monocyte chemoattractant protein-1 (MCP-1, Fig S2B), a key chemokine regulating migration and infiltration.(Singh et al., 2021) as well as IL-8.(Fig S2C).

### Hepcidin induces mitochondrial dysfunction in PASMC but not in PAEC

To investigate if responses to hepcidin treatment and associated cellular iron retention within PAEC influenced mitochondrial function, cells were given direct hepcidin treatments for 24 hours. PASMC were similarly challenged to determine variations between both cell types.

#### Morphological Changes

In PAEC, mitochondrial networks appeared to bud and became disrupted following hepcidin treatment, but no fractionation was evident as shown (Fig 3A). By contrast, mitochondria in PASMCs exposed to hepcidin demonstrated obvious changes in morphology as strikingly defined networks became fractionated (Fig 3B). In addition, PASMC mitochondria showed no budding in contrast to PAECs but did appear less uniform in their arrangement following hepcidin treatment (Fig 3B). Because Mitotracker Orange (CMTMros) enters mitochondria based on mitochondrial membrane potential (MMP) (Elmore et al., 2004), fluorescence can infer an immediate measurement of MMP, however, this is not a reliable metric over time (Kholmukhamedov et al., 2013). Whilst no change to fluorescence was revealed in hepcidin treated PAEC (Fig 3C), it was significantly reduced in similarly treated PASMC (Fig 3D) indicative of a reduction in MMP. This observation was supported by subsequent seahorse experiments.

**Figure 3.**
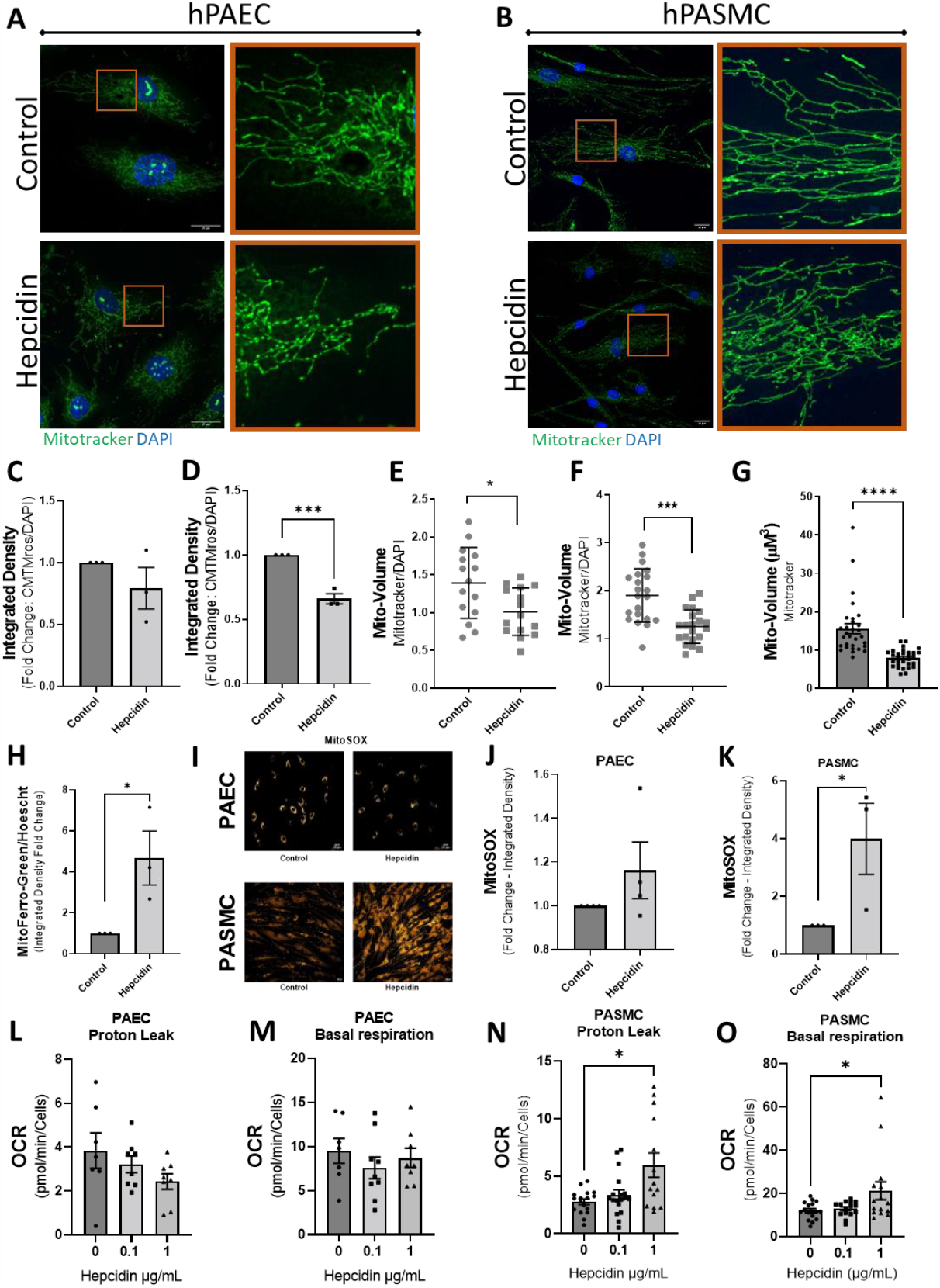
Hepcidin treatment produces mitochondrial effects in hPAEC and hPASMC. hPAEC **(A)** and hPASMC **(B)** were stained with Mitotracker (green) and DAPI (blue) following treatment with or without hepcidin. Orange boxes show magnified region. Scale bars, 20µm. **(C)** Integrated density of hPAEC Mitotracker was quantified and represented as average fold change relative lo control for multiple eels per condition (n = 3). **(D)** Integrated density of hPASMC Mitotracker was quantified and represented as fold change relative lo control as with C (n=3). **(E)** Mitochondrial volume of hPAEC eels following treatment with or without hepcidin. Mitochondrial volume normalized to nuclear (DAPI) volume(n = 15, 3 experimental repeats). **(F)** Mitochondrial volume of hPASMC following treatment with or without hepcidin. Mitochondrial volume normalized to nuclear (DAPI) volume (n= 24, 3 experimental repeats). **(G)** Mitochondrial volume (µM^3^), determined using mitotracker for PASMC following treatment with or without hepcidin (n=30, 3 experimental repeats). **(H)** Fold change of integrated density for MitoFerro green normalized to Hoescht for PASMC (mean ± SEM; n=3). (Q Representative images of PAEC and PASMC treated with or without hepcidin and stained with MitoSOX. **(J)** Quantification of MitoSOX integrated density for PAEC shown as fold change hepcidin vs control (mean± SEM; n=4). **(K)** Quantification of MitoSOX integrated density for PAEC shown as fold change hepcidin vs control (mean ± SEM; n=3). (L) Proton leak and **(M)** basal respiration from PAEC treated with hepcidin. Oxygen consumption rate (OCR) was determined by Seahorse XF analyser mito stress test (n= 7-10). **(N)** Proton leak and (0) basal respiration from PASMC treated with hepcidin (n = 15-17). students T-test was perfomed or ANOVA with bonferonni post-hoc test for L-0.Mean ± SEM for all bar charts.-p <0.001;^*^p <0.05

Moreover, PAEC showed reduced (∼25%) mitochondrial volume per cell in response to hepcidin treatment (Fig 3E) whereas in PASMC, a much more obvious significant reduction in per cell mitochondrial volume following 24 hr hepcidin of ∼50% was seen (Fig 3F). For these analyses mitochondrial volume was normalised per cell against nuclear volume to account for variations in cell size. Importantly, in this regard, nuclear volume showed no change in response to hepcidin treatment at 24hrs in PASMC (Fig S3A). Furthermore, the change in mitochondrial volume observed were confirmed by analysing mitochondrial volume using mitochondrial marker TOM20, which revealed significant reductions in average mitochondrial volume in PASMC following 24 hours hepcidin (Fig S3B).

#### Fragmentation of networks were observed in PASMC

This was quantified as a change in average mitochondrial volume which revealed smaller detectable units. A larger average reveals the continuation of a network and smaller average is indicative of a broken, fragmented network. Treatment of PASMC with hepcidin for 24 hr revealed a significant reduction in average mitochondrial volume (Fig 3G).

#### Mitochondrial iron retention and ROS production

Changes to mitochondrial iron levels were determined by use of Mito-FerroGreen, a mitochondria-specific iron probe that becomes fluorescent when reacting with labile mitochondrial Fe^2+^ was used (Hirayama et al., 2018). In PAECs incubated with hepcidin, increases in mitochondrial specific iron as determined by increases in mito-ferrogreen staining (Fig S3C), was expressed as integrated density (Fig 3H). It proved technically challenging to use this technique in PASMCs. However, we have previously reported PASMC iron accumulation in response to hepcidin challenge although mitochondrial specific iron determinations were not undertaken in that study (Ramakrishnan et al., 2018).

To further investigate cell specific mitochondrial responses, assessment of mitochondrial ROS production was undertaken using the mitochondrial specific dye, MitoSOX, which becomes fluorescent upon superoxide-mediated oxidation. Immunoflourescence showed no noticeable change in MitoSOX staining following hepcidin treatment in PAECs but an obvious response in PASMCs (Fig 3I), this was quantified for PAEC (Fig 3J) and PASMC (Fig 3K).

Mitochondrial function was further investigated through seahorse assay. Proton leak measures the number of protons entering the extracellular space, indicating changes in membrane potential and membrane function. Increases in proton leak indicate mitochondrial damage (Cheng et al., 2017). Basal respiration describes the oxygen consumption used to meet cellular AP demand resulting from mitochondrial proton leak. Changes indicate energetic demand of cells under baseline conditions. Hepcidin treatment revealed little change in dynamics of PAEC as measured by seahorse, with a slight decrease in proton leak (Fig 3L) and a very slight decrease in basal respiration (Fig 3M). In contrast, there was a significant increase in proton leak (Fig 3N) and basal respiration (Fig 3O) for hPASMC treated with 1 µg/mL hepcidin.

### Conditioned media from hepcidin treated PAEC induces migration, proliferation and mitochondrial responses in PASMC

#### Ferroportin expression

Confocal imaging of ferroportin in PASMCs incubated for 24 hr with conditioned media obtained from PAECs following 24 hours of treatment with hepcidin (1 µg/ml) resulted in a complete loss of ferroportin staining in these cells by confocal imaging (Fig 4A), by contrast media from untreated PAECs or hepcidin in media only controls was unable to affect ferroportin staining in these cells (Fig 4A). These results suggest an endothelial derived component, most likely hepcidin, (see Fig 2D) is responsible for this decisive inhibitory response. Given these findings, relevant functional responses in relation to said conditioned media were then assessed in PASMCs.

**Figure 4:**
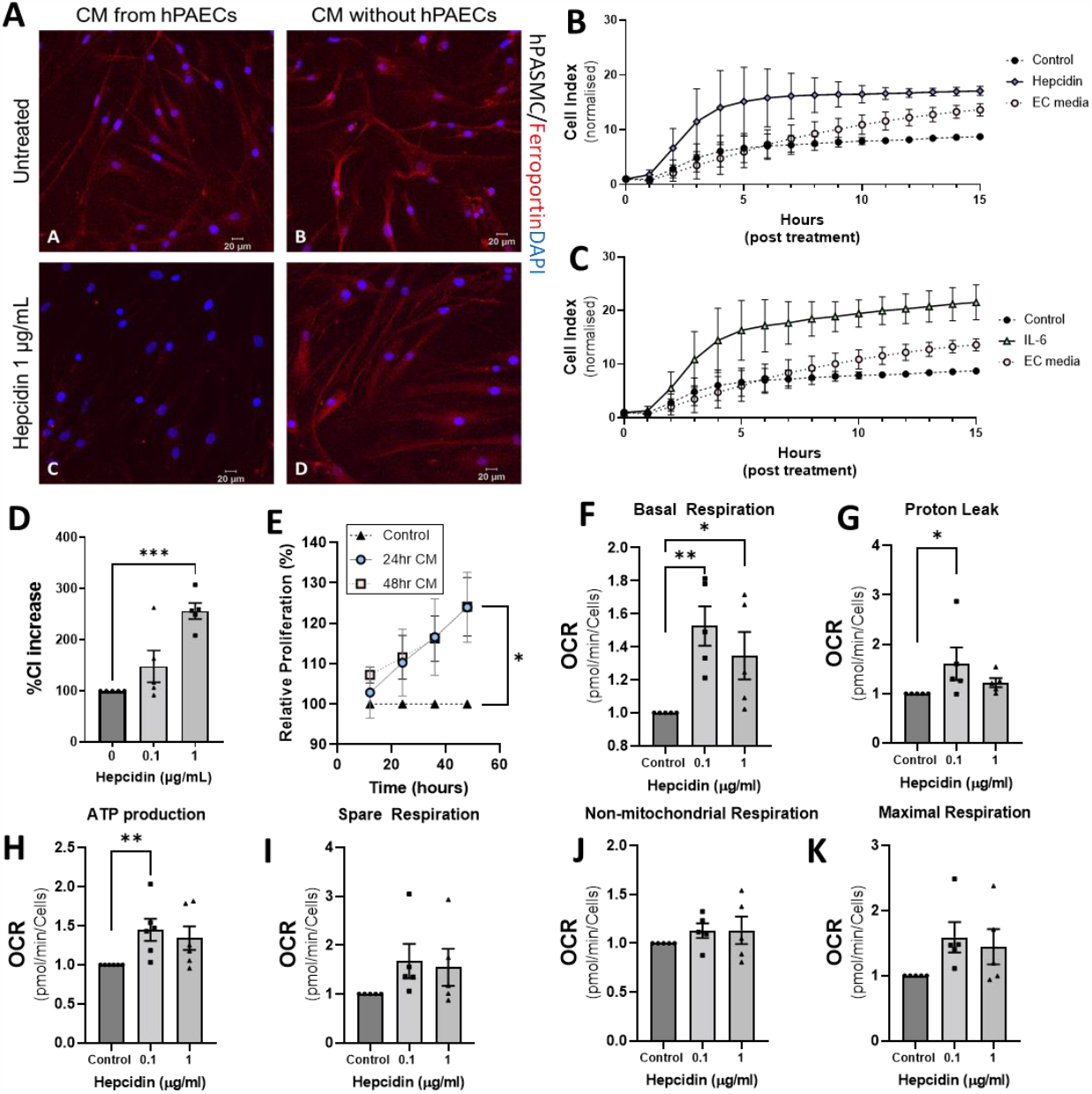
Effects of hPAEC conditioned media treated with hepcidin on hPASMC. Immunoflourescent images of hPASMC expression ferroportin after 24 hours exposure to hPAEC conditioned media **(A)**. Ferroportin (red) and DAPI (blue). Changes in pulmonary artery smooth muscle cell migration in hepcidin-treated conditioned media **(B)** and IL-6 treated conditioned media **(C)** or endothelial cell (EC) media alone measured by exCELLigence assay. **(D)** Quanitifation of hPASMC migration at 8 hours with treatment of conditioned media treated with hepcidin. **(E)** Initial hPASMC proliferation in response to 24 and 48 hr conditioned media with or without hepcidin addition, measured by exCELLigence assay. **(F-K)** hPASMC mitochondria metabolism changes after 24 hours treatment with hPAECs conditioned media. Oxygen consumption rate (OCR) was determined by Seahorse XF analyser mito stress test. Data shown are mean ±SEM n=6. Kruskal-Wallis test followed by Dunn’s post hoc test was performed for all but E, where two-way ANOVA with Tukey post hoc analysis was performed ^*^p<0.05; ^**^p<0.01, p<0.001.

#### PASMC migration

The The xCELLigence Real Time Cell Analyser (RTCA) system was used to determine any effects of PAEC conditioned media on PASMC migration. Results showed that 24 h conditioned media either from IL-6 or hepcidin challenged PAECS caused significant PASMC migration (Fig 4B and C). This was significant by 8 hours of exposure when compared to untreated PAEC conditioned media controls for 1 ng/mL of IL-6 (Fig S4A) and 1 µg/mL hepcidin (Fig 4B,D and S4A). Conditioned media with 0.1 and 1µg/mL revealed concentration specific increases in this migratory response in PASMC treated with conditioned media at 8 hours (Fig 4D). Additional studies using direct hepcidin treatments at the same concentrations in media alone did not cause migration, so excluding the possibility that these agents were acting as directly proliferative agents (Fig S4B) and confirming that the migratory responses observed were PAEC mediated. Furthermore, PAEC media alone did not induce this migratory response (Fig S4A) and it was hepcidin treatment which induced this response. PASMC migration was observed to be increased to a significantly greater extent in 24 hr conditioned media than 48 hr (Fig S4D).

#### PASMC proliferation

The xCELLigence Real Time Cell Analyser (RTCA) system was used to determine any effects of hPAEC conditioned media on PASMC proliferation. Cell index measurements revealed significantly increased PASMC proliferation as an initial rate of proliferation in response to treatment with 48 hour PAEC hepcidin conditioned media but not with 24 hour conditioned media (Fig S4D). In addition, as cell index measurements are made by the xCelligence RTCA system continuously in real time over a period of 48 hours assessment of total proliferation was also undertaken. Analysis of data was limited to 4 time points (12, 24, 36 and 48 hours). Overall, conditioned media (CM) from hepcidin treated PAECs significantly increased the total proliferation of PASMCs after 48 hours (Fig 4E). CM from 24 hour treated PAECs increased total 48 hour proliferation by PASMCs by 24% (95% CI 1.89 – 46.1, P<0.05), and CM from 48 hour treated PAECs by 24% (95% CI 0.256-47.9, P<0.05). No significant effect of conditioned media was seen at 12, 24, or 36 hour time points but there was an observable trend of proliferation increasing with time (Fig 4E).

To further investigate cellular state in PASMC in response to direct treatment with hepcidin or IL-6, cell cycle was investigated with flow cytometry. Hepcidin at 1 µg/mL induced a significant reduction in cells in G0/G1 and increases in the growth phases, S and G2/M (FigS4E). A similar effect was shown with 10ng/mL IL-6 at 24 hours with increases in cells in S and G2/M (Fig S4F).

Proliferation of PAEC was also assessed at 24 hours for direct treatment of hepcidin and IL-6 with BrdU assay. At 24 hours hepcidin challenge revealed no change in proliferation vs control (Fig S4E) whereas IL-6 treatment revealed a significant increase in proliferation relative to control (Fig S4F). Cell cycle analysis of PAEC following 24 hours treatment with hepcidin or IL-6 revealed no changes compared to control cells (Fig S4I and J).

PAEC viability was investigated using MTS assay. Direct treatment of hepcidin revealed a slight insignificant increase in viability (Fig S4X) whereas 10 ng/ml of IL-6 treatment increased viability significantly compared to control (Fig S4X). Further to this, cell death at varying concentrations up to 1 µg/mL hepcidin and 10 ng/mL IL-6 was investigated with alarmar blue assay and no significant changes were observed (Fig S4M and O).

#### Mitochondrial respiration

Given the dramatic effects of conditioned media on ferroportin expression seen in Fig 4A and the studies presented in Fig 3 which demonstrated considerable levels of mitochondrial dysfunction in PASMCs treated directly with further investigations of mitochondrial respiration were undertaken. PAEC treated with hepcidin produced significant changes in basal respiration (Fig 4F), proton leak (Fig 4G) and ATP production (Fig 4H). Hepcidin treatment also produced increases in spare respiration (Fig 4I), Non-mitochondrial respiration (Fig 4J) and maximal respiration (Fig 4K). Further to this, the effects of 48 hour conditioned media upon PASMC mitochondrial respiration was also investigated. In contrast to the effects of 24 hour conditioned media, maximal respiration and spare respiration were both significantly reduced (Fig S5A and B). Basal respiration, proton leak and non-mitochondrial respiration also all showed relative reduction compared to control (Fig S5C, D and E). Together these data present a considerable change to PASMC metabolism in response to hepcidin treated PAEC conditioned media treatment.

Given that hepcidin has previously been shown to increase proliferation in PASMC (Ramakrishnan et al., 2018) and that studies presented in figure 2 and described above indicate that hepcidin treatment of PAECs caused greatly augmented cellular release of hepcidin, it seems plausible to suggest a causative link between PAEC hepcidin release and PASMC proliferation, outlined in the graphical abstract.

## Discussion

The studies presented here indicate a key regulatory function for expression and modulation of the hepcidin-ferroportin axis within cells of the pulmonary vasculature. Moreover, crosstalk between PAECs and PASMCs indicates an important paracrine role for hepcidin and other associated molecules in this setting.

### Modulation of the hepcidin-ferroportin axis in PAEC and PASMC

There are few studies demonstrating the presence of ferroportin in endothelial cells although it has been described in brain, cardiac and retinal cells (Wu et al., 2004, Lakhal-Littleton et al., 2015, Baumann et al., 2019). The research presented here (fig 1) clearly demonstrates the presence of ferroportin in PAECs and regulation in response to hepcidin or IL-6 treatment. Furthermore, we observed upregulation of hepcidin mRNA and hepcidin in the media of PAECs treated with IL-6. These findings confirm and expand preliminary observations in PAECs (Toe et al., 2022) and further demonstrate modulation, confirming observations seen in PASMCs (Ramakrishnan et al., 2018); with evidence of increased proliferation (Fig 4E), although limited in nature.

IL-6 has been implicated as a potential mediator in PAH (Savai et al., 2012, Pullamsetti et al., 2018) but a recent clinical trial using the IL-6 receptor antagonist, tocilizumab, failed to report any benefit in PAH patients (Toshner et al., 2022). In contrast, we present a potential role for IL-6 modulation of the hepcidin-ferroportin axis and proliferation in PAECs. However, IL-6 is not the exclusive positive regulator of this axis; interferon alpha (Ichiki et al., 2014) and IL-1 (Lee et al., 2005) amongst others, also up-regulate hepcidin production in cells types more traditionally associated with iron homeostasis; as yet no evaluations have been undertaken in PAECS.

Our research demonstrates IL-6 gene transcription and release from PAECs in response to hepcidin, indicating potential localised regulation of iron homeostasis in these cells. Unexpectedly, it was observed that hepcidin treatment significantly enhanced hepcidin (HAMP) mRNA production and hepcidin release from PAECs, supported by observed modulation by siRNA knockdown studies of HAMP. Comparative media studies allowed us to demonstrate hepcidin production by PAECs was not an artefact of treatment dose and hepcidin has a short half-life in biological matrices. (Ruchala and Nemeth, 2014). To determine that loss of (dosage) hepcidin was not a binding effect which could be reversed by conditional changes in media over time, buffered saline containing hepcidin, again, there was a rapid loss in measured hepcidin over the time course of the experiment to undetectable levels (data not shown). This Study clearly indicates the involvement of PAECs in hepcidin production and release, demonstrating the potential for localised regulation of the hepcidin-ferroportin axis in these cells.

### Hepcidin modulates mitochondrial and respiratory function in PAEC and PASMC

Because modulation of the hepcidin-ferroportin axis likely impacts cellular iron resources and fluxes impacting mitochondrial function, we investigated mitochondrial function in both PASMC and PAEC following hepcidin treatment. Confocal imaging and analysis of mitochondrial networks showed distinct changes in mitochondrial morphology and membrane potential for both cell types post hepcidin treatment; however, it was apparent PAEC mitochondria were more resistant to hepcidin treatment than PASMC mitochondria. Additionally, mitochondrial reactive species production was apparent in PASMCs but not replicated in PAECs.

A role for mitochondrial iron-loading is well established in health and disease (Issitt et al., 2019, Bi et al., 2021, Sawicki et al., 2023). Importantly, mitochondrial responses described here, are similar to responses observed in mesenchymal stromal cells where iron overload resulted in mitochondrial fragmentation in a ROS dependant manner (Zheng et al., 2018).

Increased mitochondrial iron retention was observed in PAEC in response to hepcidin. Technical issues with mitoferro-green prevented comparative examination in PASMCs but we have previously reported iron loading in PASMC in response to hepcidin (Ramakrishnan et al., 2018). Hepcidin treatment produced observable mitochondrial respiratory dysfunction assessed by seahorse assay, presumably through iron retention, in a more significant manner in PASMC than PAEC. Changes in respiration may suggest a shift to glycolytic metabolism, previously reported to be linked to aspects of disease pathology in PAH (Archer, 2017). Together, our reported mitochondrial studies clearly demonstrated that mitochondria of PAECs are much more by modulation by the hepcidin-ferroportin axis than PASMCs.

### Media from PAECs treated with hepcidin induces proliferative, migratory and respiratory changes in PASMCs

Our data indicates that PAECs produce and release significant amounts of hepcidin in response to treatment with hepcidin. To explore the potential relationship of the hepcidin-ferroportin axis between PAECs and PASMCs, conditioned media from PAECs treated with hepcidin was exposed to PASMCs. A striking, almost complete, loss of ferroportin staining was apparent in PASMCs treated this way when compared to untreated PAEC conditioned media. Furthermore, media conditioned without PAECs over the same incubation time frame was unable to cause loss of ferroportin staining either alone or with hepcidin included. These results strongly indicate the necessity for PAEC involvement and are indicative of endogenous hepcidin production by PAECs. In addition, the conditioned media was able to induce significant migratory response in PASMCs and a significant proliferative response when media was conditioned for 48 hrs (see Fig S4D).

Pronounced changes in mitochondrial respiration were observed with 24 and 48 hour conditioned media. We were unable to find an intervention to specifically target hepcidin in conditioned media, but PAEC conditioned media controls lacking hepcidin were unable to replicate findings seen with PAEC hepcidin conditioned media, which strongly suggests that hepcidin produced by PAECs is the active component facilitating mitochondrial dysfunction and PASMC proliferation.

Indeed, mitochondria are known to be involved in aspects of cell fate including proliferation (Antico Arciuch et al., 2012, Kirova et al., 2022). As for the rapid and extensive PASMC migratory responses observed in PASMC exposed to PAEC hepcidin conditioned media, it is not obvious how hepcidin would facilitate this response as there is no literature to our knowledge indicating that hepcidin has chemotactic properties. Moreover, hepcidin alone in media was unable to cause migration of PASMCS (Fig S4A) reinforcing this notion. However, significantly elevated levels of cytokines were also seen in such conditioned media (Fig S2B and C) may well offer a plausible explanation for the migratory responses observed. MCP-1 and IL-8 are all known to possess chemoattractant properties (Singh et al., 2021, Cesta et al., 2021) including with relevance for pulmonary vascular migratory responses in PAH (Rabinovitch et al., 2014, Soon et al., 2010, Arends et al., 2013). Based on our findings the proposed mechanism for PAH progression at the level of the pulmonary vasculature is best illustrated in the graphical abstract.

### Summary

These studies have demonstrated the presence of and effective modulation of a hepcidin-ferroportin iron regulatory axis in PAECs. It is also apparent that PAECs are more tolerant of exposure to hepcidin as evidenced by limited changes to mitochondrial morphology, ROS production and respiration in comparison to that seen with PASMCs. The pronounced effects of conditioned media from PAECs on hPASMC function, including loss of ferroportin expression, altered mitochondrial respiration and ultimately enhanced proliferation, strongly implicate hepcidin produced from PAECs as the driver of these responses. It is likely that enhanced iron retention in PASMCs due to loss of ferroportin mediated iron export, stimulated proliferation, both via mitochondrial related responses in addition to more classically recognised mechanisms.

Further evaluation of the hepcidin-ferroportin axis under low oxygen tensions, with specific sensitive measures of cellular iron pools and in relevant in vivo models of PAH may well offer additional insight and strengthen the case for therapeutic intervention to target this axis in patients with PAH.

## Abbreviations

(PASMCs): Human Pulmonary artery smooth muscle cells
(PAECs): Human pulmonary artery endothelial cells
(PAH): pulmonary artery hypertension
IL-6: interleukin-6

## Acknowledgements

The authors would like to acknowledge the considerable impact the late Dr Latha Ramakrishnan had upon the work presented here and the wider field of iron regulation and pulmonary function.

We would also like to thank the Dinosaur trust who provided the eXcelligence system.

## References

Antico Arciuch, V. G., Elguero, M. E., Poderoso, J. J. & Carreras, M. C. 2012. Mitochondrial regulation of cell cycle and proliferation. Antioxid Redox Signal, 16, 1150–80.

Archer, S. L. 2017. Pyruvate Kinase and Warburg Metabolism in Pulmonary Arterial Hypertension: Uncoupled Glycolysis and the Cancer-Like Phenotype of Pulmonary Arterial Hypertension. Circulation, 136, 2486–2490.

Arends, S. J., Damoiseaux, J. G., Duijvestijn, A. M., Debrus-Palmans, L., Boomars, K. A., Brunner-La Rocca, H. P., Cohen Tervaert, J. W. & Van Paassen, P. 2013. Functional implications of IgG anti-endothelial cell antibodies in pulmonary arterial hypertension. Autoimmunity, 46, 463–70.

Baumann, B. H., Shu, W., Song, Y., Simpson, E. M., Lakhal-Littleton, S. & Dunaief, J. L. 2019. Ferroportin-mediated iron export from vascular endothelial cells in retina and brain. Exp Eye Res, 187, 107728.

Bi, Y., Ajoolabady, A., Demillard, L. J., Yu, W., Hilaire, M. L., Zhang, Y. & Ren, J. 2021. Dysregulation of iron metabolism in cardiovascular diseases: From iron deficiency to iron overload. Biochem Pharmacol, 190, 114661.

Cesta, M. C., Zippoli, M., Marsiglia, C., Gavioli, E. M., Mantelli, F., Allegretti, M. & Balk, R. A. 2021. The Role of Interleukin-8 in Lung Inflammation and Injury: Implications for the Management of COVID-19 and Hyperinflammatory Acute Respiratory Distress Syndrome. Front Pharmacol, 12, 808797.

Cheng, J., Nanayakkara, G., Shao, Y., Cueto, R., Wang, L., Yang, W. Y., Tian, Y., Wang, H. & Yang, X. 2017. Mitochondrial Proton Leak Plays a Critical Role in Pathogenesis of Cardiovascular Diseases. Adv Exp Med Biol, 982, 359–370.

Condon, D. F., Nickel, N. P., Anderson, R., Mirza, S. & De Jesus Perez, V. A. 2019. The 6th World Symposium on Pulmonary Hypertension: what’s old is new. F1000Research, 8, F1000 Faculty Rev–888.

Cotruvo, J. A. & Stubbe, J. 2011. Class I ribonucleotide reductases: metallocofactor assembly and repair in vitro and in vivo. Annu Rev Biochem, 80, 733–67.

Dannewitz Prosseda, S., Ali, M. K. & Spiekerkoetter, E. 2020. Novel Advances in Modifying BMPR2 Signaling in PAH. Genes (Basel),

Drakesmith, H., Nemeth, E. & Ganz, T. 2015. Ironing out Ferroportin. Cell Metab, 22, 777–87.

Elmore, S. P., Nishimura, Y., Qian, T., Herman, B. & Lemasters, J. J. 2004. Discrimination of depolarized from polarized mitochondria by confocal fluorescence resonance energy transfer. Archives of Biochemistry and Biophysics, 422, 145–152.

Finberg, K. E. 2013. Regulation of systemic iron homeostasis. Curr Opin Hematol, 20, 208–14.

Ginzburg, Y. Z. 2019. Hepcidin-ferroportin axis in health and disease. Vitam Horm, 110, 17–45.

Graf, S., Haimel, M., Bleda, M., Hadinnapola, C., Southgate, L., Li, W., Hodgson, J., Liu, B., Salmon, R. M., Southwood, M., Machado, R. D., Martin, J. M., Treacy, C. M., Yates, K., Daugherty, L. C., Shamardina, O., Whitehorn, D., Holden, S., Aldred, M., Bogaard, H. J., Church, C., Coghlan, G., Condliffe, R., Corris, P. A., Danesino, C., Eyries, M., Gall, H., Ghio, S., Ghofrani, H. A., Gibbs, J. S. R., Girerd, B., Houweling, A. C., Howard, L., Humbert, M., Kiely, D. G., Kovacs, G., Mackenzie Ross, R. V., Moledina, S., Montani, D., Newnham, M., Olschewski, A., Olschewski, H., Peacock, A. J., Pepke-Zaba, J., Prokopenko, I., Rhodes, C. J., Scelsi, L., Seeger, W., Soubrier, F., Stein, D. F., Suntharalingam, J., Swietlik, E. M., Toshner, M. R., Van Heel, D. A., Vonk Noordegraaf, A., Waisfisz, Q., Wharton, J., Wort, S. J., Ouwehand, W. H., Soranzo, N., Lawrie, A., Upton, P. D., Wilkins, M. R., Trembath, R. C. & Morrell, N. W. 2018. Identification of rare sequence variation underlying heritable pulmonary arterial hypertension. Nat Commun, 9, 1416.

Hirayama, T., Kadota, S., Niwa, M. & Nagasawa, H. 2018. A mitochondria-targeted fluorescent probe for selective detection of mitochondrial labile Fe(ii). Metallomics, 10, 794–801.

Ichiki, K., Ikuta, K., Addo, L., Tanaka, H., Sasaki, Y., Shimonaka, Y., Sasaki, K., Ito, S., Shindo, M., Ohtake, T., Fujiya, M., Torimoto, Y. & Kohgo, Y. 2014. Upregulation of iron regulatory hormone hepcidin by interferon alpha. J Gastroenterol Hepatol, 29, 387–94.

Issitt, T., Bosseboeuf, E., De Winter, N., Dufton, N., Gestri, G., Senatore, V., Chikh, A., Randi, A. M. & Raimondi, C. 2019. Neuropilin-1 Controls Endothelial Homeostasis by Regulating Mitochondrial Function and Iron-Dependent Oxidative Stress. iScience, 11, 205–223.

Kholmukhamedov, A., Schwartz, J. M. & Lemasters, J. J. 2013. Isolated mitochondria infusion mitigates ischemia-reperfusion injury of the liver in rats: mitotracker probes and mitochondrial membrane potential. Shock, 39, 543.

Kirova, D. G., Judasova, K., Vorhauser, J., Zerjatke, T., Leung, J. K., Glauche, I. & Mansfeld, J. 2022. A ROS-dependent mechanism promotes CDK2 phosphorylation to drive progression through S phase. Dev Cell, 57, 1712–1727 e9.

Lakhal-Littleton, S., Crosby, A., Frise, M. C., Mohammad, G., Carr, C. A., Loick, P. A. M. & Robbins, P. A. 2019. Intracellular iron deficiency in pulmonary arterial smooth muscle cells induces pulmonary arterial hypertension in mice. Proc Natl Acad Sci U S A, 116, 13122–13130.

Lakhal-Littleton, S., Wolna, M., Carr, C. A., Miller, J. J., Christian, H. C., Ball, V., Santos, A., Diaz, R., Biggs, D., Stillion, R., Holdship, P., Larner, F., Tyler, D. J., Clarke, K., Davies, B. & Robbins, P. A. 2015. Cardiac ferroportin regulates cellular iron homeostasis and is important for cardiac function. Proc Natl Acad Sci U S A, 112, 3164–9.

Lee, P., Peng, H., Gelbart, T., Wang, L. & Beutler, E. 2005. Regulation of hepcidin transcription by interleukin-1 and interleukin-6. Proc Natl Acad Sci U S A, 102, 1906–10.

Lin, H. C., Liu, S. Y., Lai, H. S. & Lai, I. R. 2013. Isolated mitochondria infusion mitigates ischemia-reperfusion injury of the liver in rats. Shock, 39, 304–10.

Machado, R. D., Southgate, L., Eichstaedt, C. A., Aldred, M. A., Austin, E. D., Best, D. H., Chung, W. K., Benjamin, N., Elliott, C. G., Eyries, M., Fischer, C., Graf, S., Hinderhofer, K., Humbert, M., Keiles, S. B., Loyd, J. E., Morrell, N. W., Newman, J. H., Soubrier, F., Trembath, R. C., Viales, R. R. & Grunig, E. 2015. Pulmonary Arterial Hypertension: A Current Perspective on Established and Emerging Molecular Genetic Defects. Hum Mutat, 36, 1113–27.

Poot, M., Zhang, Y. Z., Kramer, J. A., Wells, K. S., Jones, L. J., Hanzel, D. K., Lugade, A. G., Singer, V. L. & Haugland, R. P. 1996. Analysis of mitochondrial morphology and function with novel fixable fluorescent stains. J Histochem Cytochem, 44, 1363–72.

Pullamsetti, S. S., Seeger, W. & Savai, R. 2018. Classical IL-6 signaling: a promising therapeutic target for pulmonary arterial hypertension. J Clin Invest, 128, 1720–1723.

Rabinovitch, M. 2012. Molecular pathogenesis of pulmonary arterial hypertension. J Clin Invest, 122, 4306–13.

Rabinovitch, M., Guignabert, C., Humbert, M. & Nicolls, M. R. 2014. Inflammation and immunity in the pathogenesis of pulmonary arterial hypertension. Circ Res, 115, 165–75.

Rouault TA. Mitochondrial iron overload: causes and consequences. Curr Opin Genet Dev. 2016 Jun;38:31–37. doi: 10.1016/j.gde.2016.02.004. Epub 2016 Mar 25. PMID: 27026139; PMCID: PMC5035716.

Ramakrishnan, L., Pedersen, S. L., Toe, Q. K., West, L. E., Mumby, S., Casbolt, H., Issitt, T., Garfield, B., Lawrie, A., Wort, S. J. & Quinlan, G. J. 2018. The Hepcidin/Ferroportin axis modulates proliferation of pulmonary artery smooth muscle cells. Sci Rep, 8, 12972.

Ruchala, P. & Nemeth, E. 2014. The pathophysiology and pharmacology of hepcidin. Trends Pharmacol Sci, 35, 155–61.

Savai, R., Pullamsetti, S. S., Kolbe, J., Bieniek, E., Voswinckel, R., Fink, L., Scheed, A., Ritter, C., Dahal, B. K., Vater, A., Klussmann, S., Ghofrani, H. A., Weissmann, N., Klepetko, W., Banat, G. A., Seeger, W., Grimminger, F. & Schermuly, R. T. 2012. Immune and inflammatory cell involvement in the pathology of idiopathic pulmonary arterial hypertension. Am J Respir Crit Care Med, 186, 897–908.

Sawicki, K. T., De Jesus, A. & Ardehali, H. 2023. Iron Metabolism in Cardiovascular Disease: Physiology, Mechanisms, and Therapeutic Targets. Circ Res, 132, 379–396.

Singh, S., Anshita, D. & Ravichandiran, V. 2021. MCP-1: Function, regulation, and involvement in disease. Int Immunopharmacol, 101, 107598.

Soon, E., Holmes, A. M., Treacy, C. M., Doughty, N. J., Southgate, L., Machado, R. D., Trembath, R. C., Jennings, S., Barker, L., Nicklin, P., Walker, C., Budd, D. C., Pepke-Zaba, J. & Morrell, N. W. 2010. Elevated levels of inflammatory cytokines predict survival in idiopathic and familial pulmonary arterial hypertension. Circulation, 122, 920–7.

Soon, E., Treacy, C. M., Toshner, M. R., Mackenzie-Ross, R., Manglam, V., Busbridge, M., Sinclair-Mcgarvie, M., Arnold, J., Sheares, K. K., Morrell, N. W. & Pepke-Zaba, J. 2011. Unexplained iron deficiency in idiopathic and heritable pulmonary arterial hypertension. Thorax, 66, 326–32.

Toe, Q. K., Issitt, T., Mahomed, A., Almaghlougth, F., Bahree, I., Sturge, C., Hu, X., Panselinas, I., Burke-Gaffney, A., Wort, S. J. & Quinlan, G. J. 2022. Human pulmonary artery endothelial cells upregulate ACE2 expression in response to iron-regulatory elements: Potential implications for SARS-CoV-2 infection. Pulm Circ, 12, e12068.

Toshner, M., Church, C., Harbaum, L., Rhodes, C., Villar Moreschi, S. S., Liley, J., Jones, R., Arora, A., Batai, K., Desai, A. A., Coghlan, J. G., Gibbs, J. S. R., Gor, D., Graf, S., Harlow, L., Hernandez-Sanchez, J., Howard, L. S., Humbert, M., Karnes, J., Kiely, D. G., Kittles, R., Knightbridge, E., Lam, B., Lutz, K. A., Nichols, W. C., Pauciulo, M. W., Pepke-Zaba, J., Suntharalingam, J., Soubrier, F., Trembath, R. C., Schwantes-An, T. L., Wort, S. J., Wilkins, M. R., Gaine, S., Morrell, N. W., Corris, P. A. & Uniphy Clinical Trials, N. 2022. Mendelian randomisation and experimental medicine approaches to interleukin-6 as a drug target in pulmonary arterial hypertension. Eur Respir J, 59.

Van De Bruaene, A., Delcroix, M., Pasquet, A., De Backer, J., De Pauw, M., Naeije, R., Vachiery, J. L., Paelinck, B., Morissens, M. & Budts, W. 2011. Iron deficiency is associated with adverse outcome in Eisenmenger patients. Eur Heart J, 32, 2790–9.

Wrighting, D. M. & Andrews, N. C. 2006. Interleukin-6 induces hepcidin expression through STAT3. Blood, 108, 3204–9.

Wu, L. J., Leenders, A. G., Cooperman, S., Meyron-Holtz, E., Smith, S., Land, W., Tsai, R. Y., Berger, U. V., Sheng, Z. H. & Rouault, T. A. 2004. Expression of the iron transporter ferroportin in synaptic vesicles and the blood-brain barrier. Brain Res, 1001, 108–17.

Yu, Y., Kovacevic, Z. & Richardson, D. R. 2007. Tuning cell cycle regulation with an iron key. Cell Cycle, 6, 1982–94.

Zhang, W., Liu, B., Wang, Y., Zhang, H., He, L., Wang, P. & Dong, M. 2022. Mitochondrial dysfunction in pulmonary arterial hypertension. Front Physiol, 13, 1079989.

Zheng, Q., Zhao, Y., Guo, J., Zhao, S., Fei, C., Xiao, C., Wu, D., Wu, L., Li, X. & Chang, C. 2018. Iron overload promotes mitochondrial fragmentation in mesenchymal stromal cells from myelodysplastic syndrome patients through activation of the AMPK/MFF/Drp1 pathway. Cell Death Dis, 9, 515.

