## Supplementary Figures for "The hepcidin-ferroportin axis influences mitochondrial function, proliferation, and migration in pulmonary artery endothelial and smooth muscle cells"

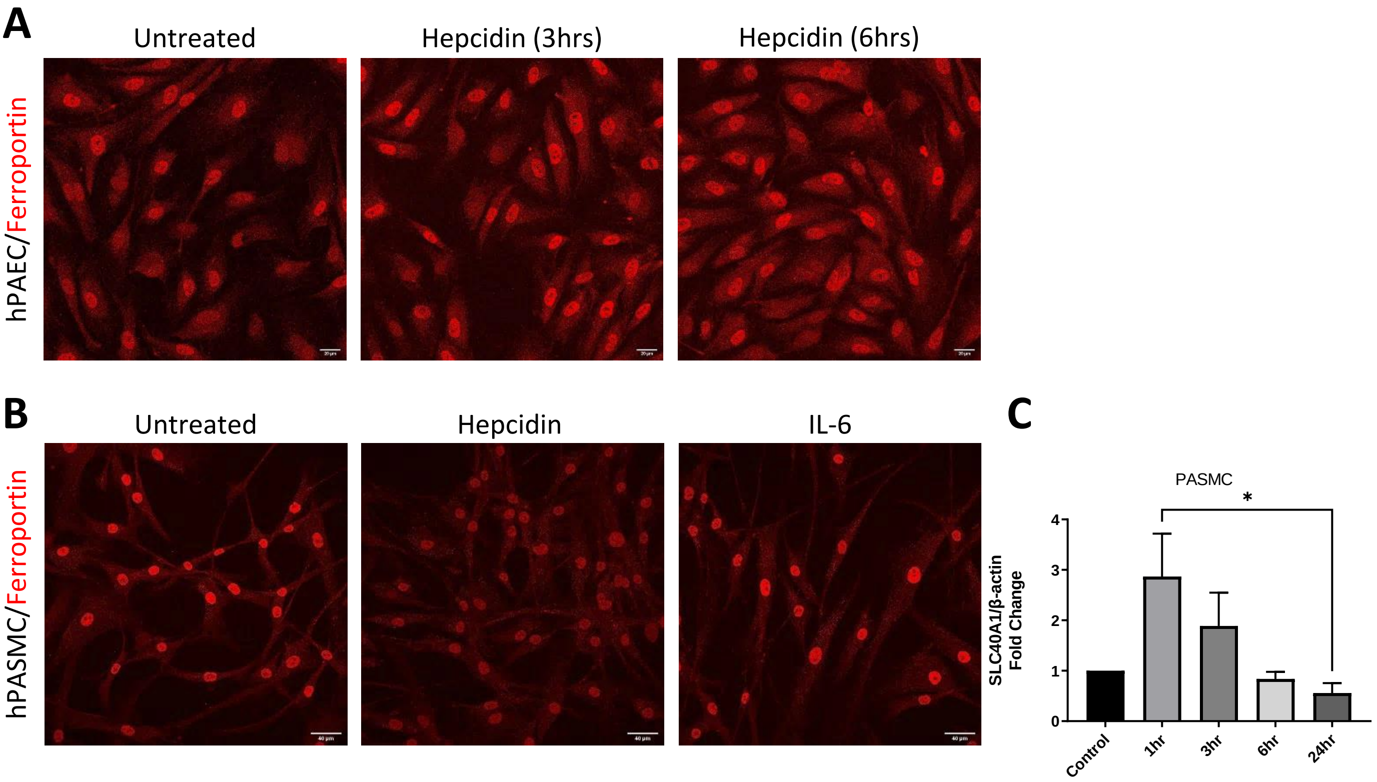


**Supplementary figure 1. Ferroportin expression and regulation in hPAEC and hPASMC. (A)** Confocal images of hPAEC grown with normal media (control) or treated with 1 μg/mL hepcidin for 3 h and 6 h and stained for Ferroportin (FPN). Scale bar 40μm. **(B)** Confocal images of hPASMC grown with normal media (control) or treated with 1 μg/mL hepcidin or 10 ng/ml IL-6 for 24 h and stained for Ferroportin (FPN). Scale bar 40μm. **(C)** Quantification of ferroportin (SLC40A1) mRNA by RT-qPCR in hPAECs treated with 1 μg/mL hepcidin, expressed as fold change of control (mean ± SEM; n = 5), ; One way ANOVA with Tukey post hoc analysis; *p < 0.05.

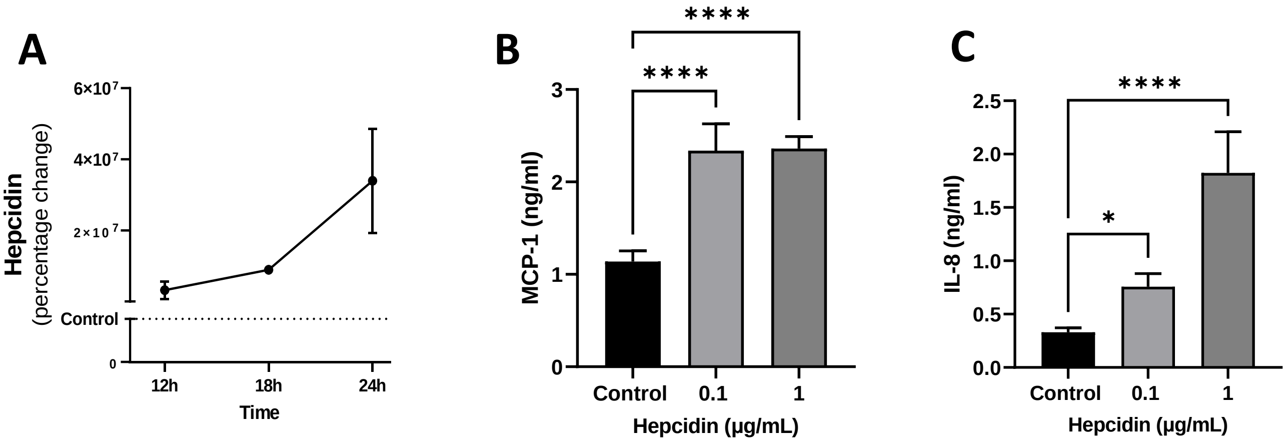


**Supplementary figure 2. Effects of hepcidin treatment on hPAEC media. (A)** Time course of hepcidin media release in response to 1 μg/mL hepcidin treatment in hPAEC measured by ELISA. Shown as percentage change relative to control (mean ± SEM; n = 5). **(B)** Monocyte chemoattractant protein-1 (MCP-1) in ng/mL determined by ELISA for media following 24 hr hepcidin treatments in hPAEC. **(C)** Quantification of ferroportin (SLC40A1) mRNA by RT-qPCR in hPAECs treated with 1 μg/mL hepcidin, expressed as fold change of control (mean ± SEM; n = 5), ; One way ANOVA with Tukey post hoc analysis for B and C; *p < 0.05 , ****p , 0.001.

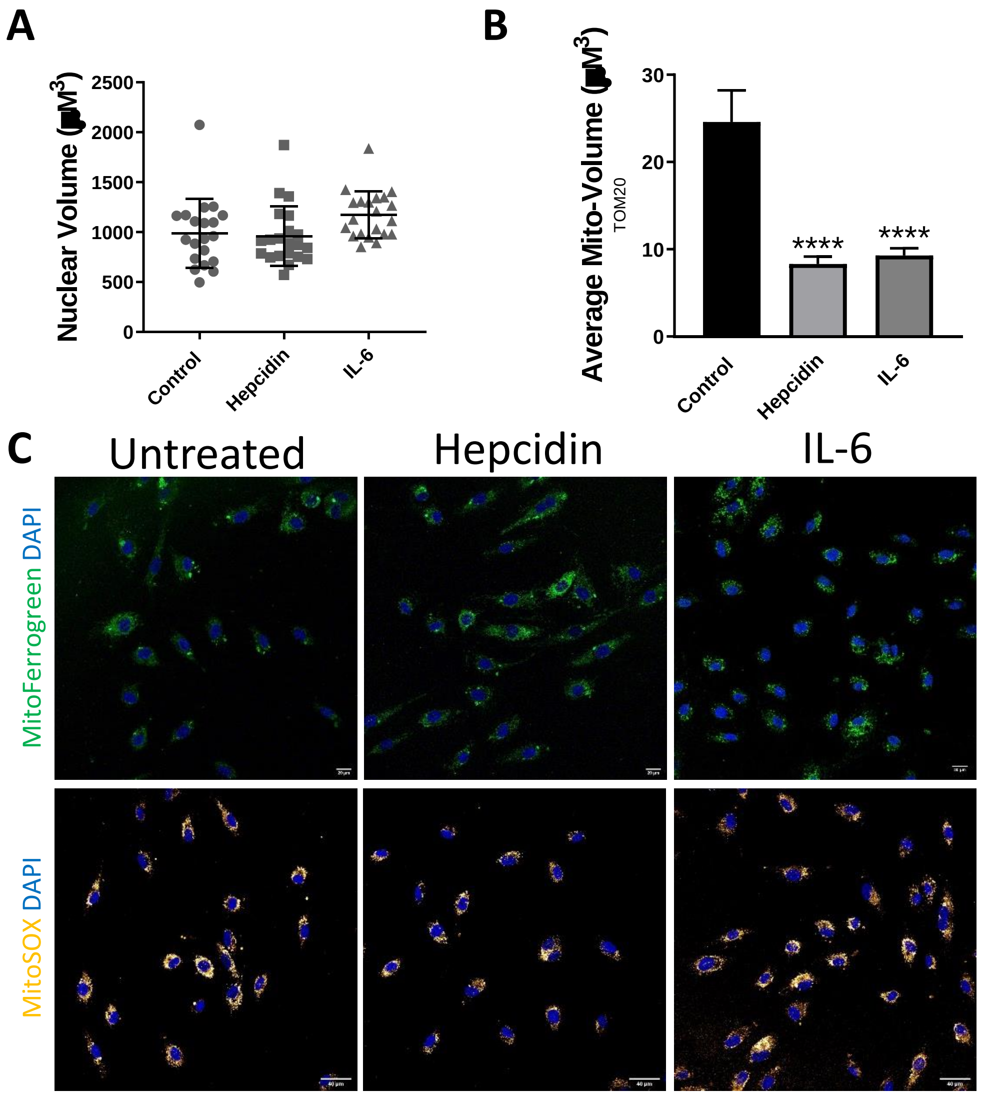


**Supplementary figure 3. Mitochondrial response to hepcidin in hPASMC and hPAEC. (A)** nuclear volume of hPASMC cells treated with hepcidin or IL-6 for 24 hours was quantified. N=3, with 21 individual cells per experiment. Error bars show standard deviation **(B)** Average mitochondrial volume for n=3, 16 cells, without normalisation, quantified using TOM20 fluorescence (mean ± SEM; n = 5), ; One way ANOVA with Tukey post hoc analysis; ****p < 0.0001. **(C)** Representative immunofluorescence for hPAEC cells treated with hepcidin or IL-6. Top panel images show mitochondrial iron content with mitoferrogreen (green) and DAPI (blue). Bottom panel images show mitochondrial reactive oxygen species using MITOSOX (yellow) and DAPI (blue).

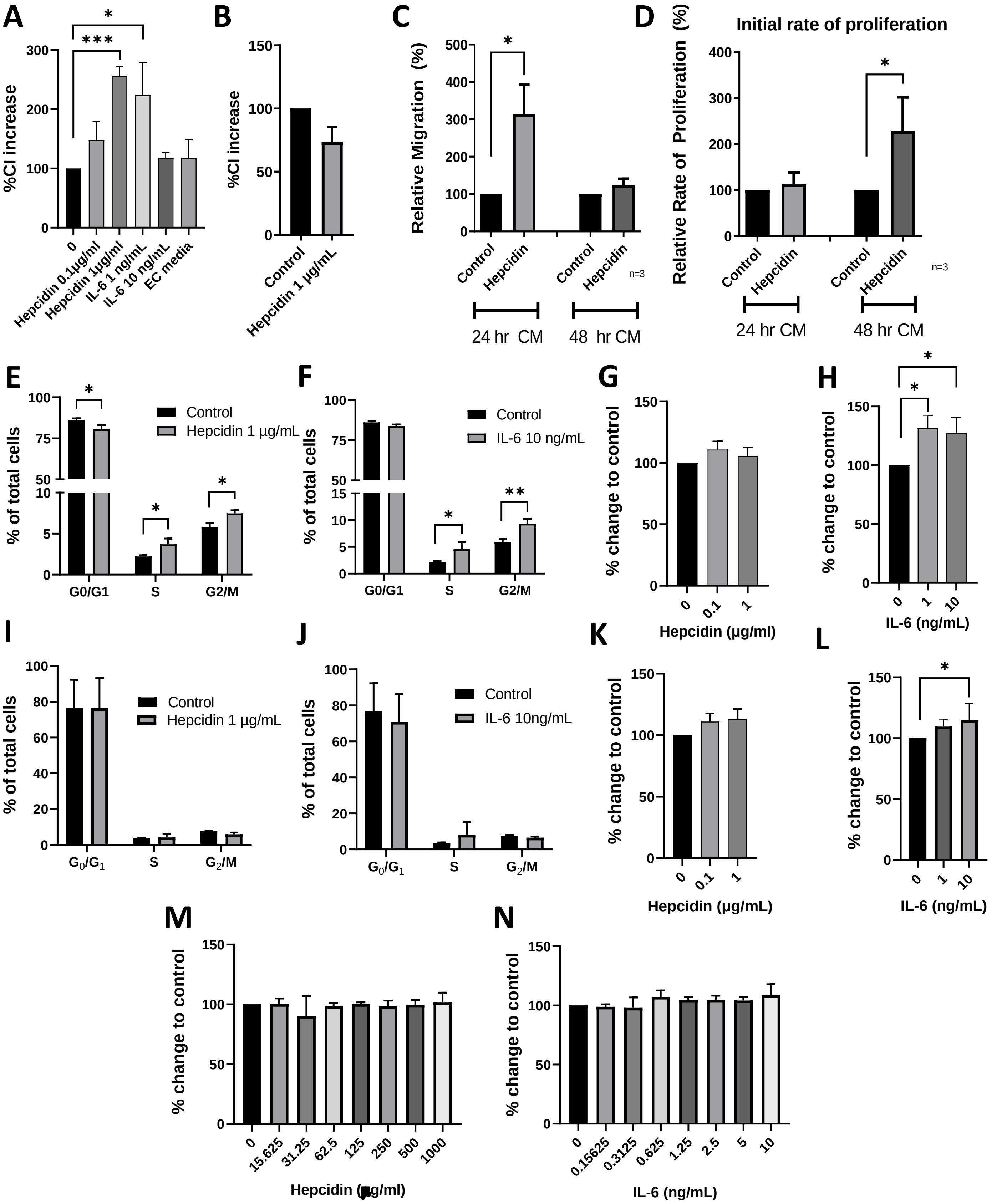


Supplementary figure 4. **Effects of conditioned media and treatments on hPASMC migration and proliferation. (A)** hPASMC migration at 8 hours for different concentrations of hepcidin and IL-6 in conditioned media. Control represents hPASMC unconditioned media. **(B)** 8 hour migration response of hPASMC treated with hepcidin containing hPAEC media without cells. **(C)** Relative migration of hPASMC in media conditioned with hPAEC for 24 or 48 hours. **(D)** Relative rate of proliferation for hPASMC treated with hPAEC conditioned media with or without hepcidin for 24 and 48 hours. **(E-F)** Cell cycle analysis of hPASMC treated with hepcidin **(E)** or IL-6 **(F)** shown as percentage of total number of cells counted. **(G- H)** hPAEC proliferation in response to direct hepcidin **(G)** and IL-6 **(H)** treatment measured by BRdU assay, shown as percentage change of control. **(I-J)** Cell cycle analysis of hPAEC treated with hepcidin **(I)** or IL-6 **(J)**, shown as percentage of total number of cells counted. **(K-L)** hPAEC viability, measured by MTS assay, for hepcidin **(K)** and IL-6 **(L)** treatments, shown as percentage change of control. **(M-N)** hPAEC cell death, measured by alamar blue assay, for hepcidin **(M)** and IL-6 **(N)** treatments, shown as percentage change of control. N=6 for A-C, G-H and K-N, n=3 for D, E-F and I-J. One-way ANOVA with Tukey post hoc analysis. *p < 0.05, **p<0.01. **p<0.001.

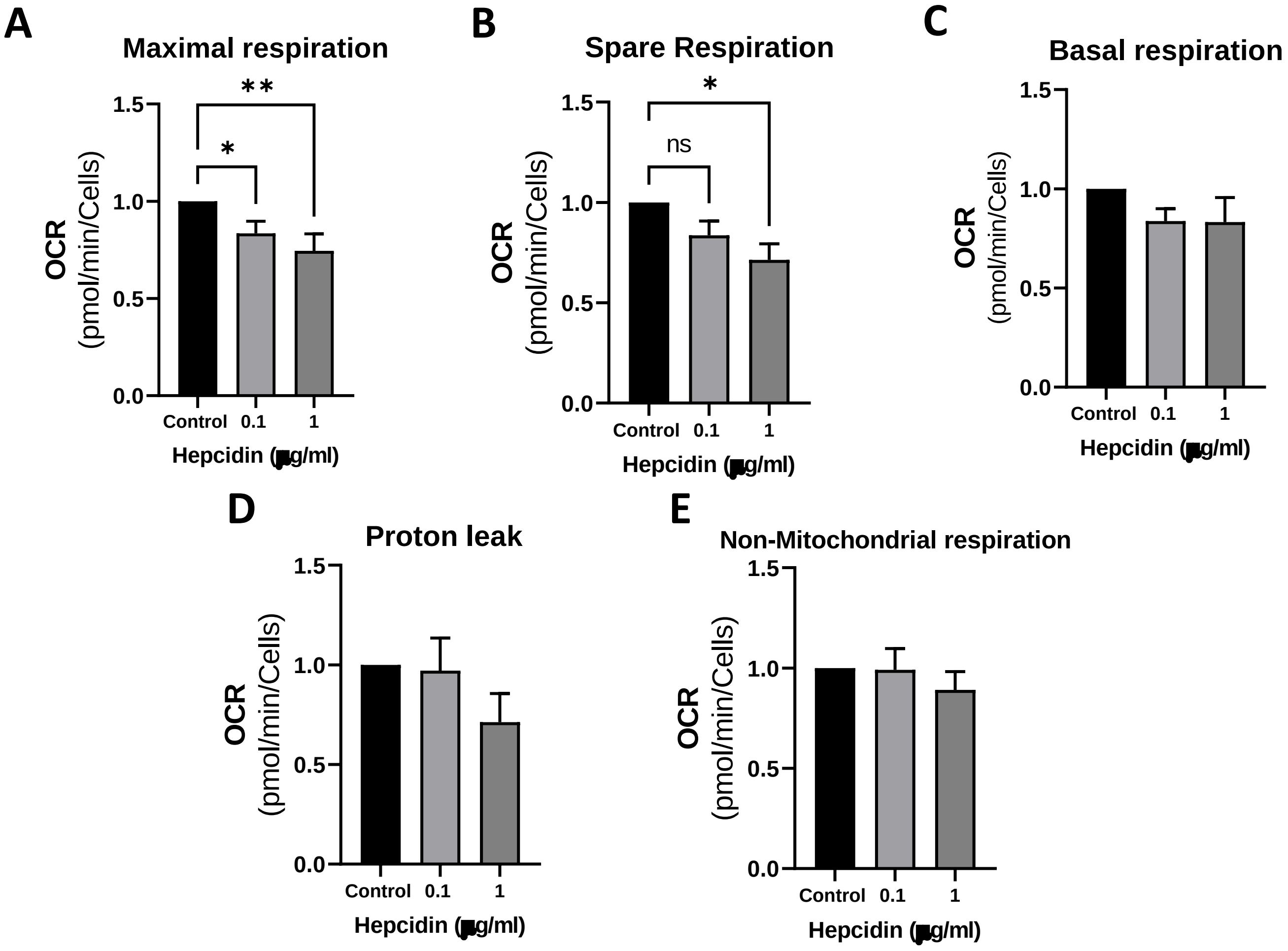


Supplementary figure 5. **Effects of 48 hour conditioned media on mitochondrial function in hPASMC.** hPASMC mitochondria metabolism changes after 48 hr treatment with hPAECs conditioned media. Oxygen consumption rate (OCR) was determined by Seahorse XF analyser mito stress test. Data shown are mean ±SEM n=6. Kruskal-Wallis test followed by Dunn's post hoc test were performed *p<0.05; **p<0.01.
